# Co-occurrence of past and present shifts current neural representations and mediates serial biases

**DOI:** 10.1101/2022.06.08.495281

**Authors:** Huihui Zhang, Huan Luo

## Abstract

The regularities of the world render an intricate interplay between past and present. Even across independent trials, current-trial perception can be automatically shifted by preceding trials, namely the ‘serial bias’. Meanwhile, the neural implementation of the spontaneous shift of present by past that operates on multiple features remains unknown. In two auditory categorization experiments with human electrophysiology recordings, we demonstrate that serial bias arises from the co-occurrence of past-trial neural reactivation and the neural encoding of current-trial features. The meeting of past and present shifts the neural representation of current-trial features and modulates serial bias behavior. Critically, past-trial features (i.e., pitch, category, motor response) that constitute an ‘event-file’ keep their respective identities in working memory and are only reactivated by the corresponding features in the current trial, giving rise to dissociated feature-specific serial biases. This ’event-file’ reactivation might constitute a fundamental mechanism for adaptive past-to-present generalizations over multiple features.

## Introduction

The regularities and recurrences of our world render the past always relevant to the present^1^. Indeed, perceptual experience and decision-making at any moment are constantly intertwined with previous information and likewise impact the future^2^, a characteristic allowing for perceptual stability and adaptive optimization. The past-to-present influences could occur automatically and span a relatively long time scale. For example, the perceived feature in the current trial (e.g., location, orientation, category) tends to be systematically shifted by that in the previous trial, even though trials are independent of each other and several seconds apart, namely the “serial bias effect” ^3–11^.

For serial bias to occur, previous trial information should leave traces in working memory (WM). Interestingly, recent studies suggest that although memories could be retained in an ‘activity-silent’ state^12–18^, memory operations still rely on the activations of the neural representations^19, 20^.

Therefore, for the past to interact with and further modify the present, previous features not only need to be retained in WM, actively or silently, but also should encounter the current features over time, that is, to be activated at the same time. A recent study demonstrated the brief reappearance of previous information during the inter-trial interval before the stimulus, supporting an integration between activity-based and activity-silent WM that contributes to the serial bias^21^. Meanwhile, the direct neural evidence for the past interacting with and altering the processing of current inputs is still lacking.

Furthermore, every perceptual decision is not simple but rich and manifold, encompassing multitudes of features at various levels, e.g., physical properties, abstract categories, response actions, etc., together constituting an “event-file”^22^. In fact, different features are associated with different serial bias directions, being attracted to or repulsed from previous ones, even within the same task^5, 7, 8, 23, 24^. Therefore, features should be compartmentalized in WM to keep their identities throughout the event-file updating across consecutive trials, in turn influencing the future in a feature-specific way.

Here we aim to understand the dynamic neural mechanisms of how the previous multi-feature event-file confronts and modifies the present, from which the feature-specific serial bias arises. In two experiments, Human subjects performed an auditory categorization task with their brain activities recorded using electroencephalography (EEG). First, behavioral results exhibit concurrent component-specific serial biases, i.e., repulsive for tone pitch and motor response, and attractive for category report. Importantly, the neural representations of features in the previous trial – pitch, category, and motor response – are reactivated and emerge simultaneously as the corresponding features in the present trial. Most crucially, the current neural representations exhibit behaviorally-congruent shift, i.e., being attracted or repulsed from the past in a feature-specific way, which further correlates to serial bias behavior.

Taken together, the past is intimately mingled with the present via an “event-file” reactivation mechanism, whereby previous information lingering in WM is triggered by the corresponding feature occurrence so that their co-emergence in time leads to the serial bias.

## Results

### Task paradigm and feature-specific behavioral serial bias

In Experiment 1, thirty human subjects performed an auditory pitch categorization task with their 64-channel EEG activities recorded, by pressing buttons to indicate the category (“high pitch” or “low pitch”) of a pure tone embedded in a sustained white noise (Fig. 1A). The pitch of the tone stimulus was pseudo-randomly selected from 5 fixed frequencies between 180 Hz and 360 Hz (f1, f2, f3, f4, f5), which were individualized to normalize task difficulty across subjects (see details in Methods). Moreover, to dissociate category report and motor response, a response cue frame appeared after the tone stimulus, based on which subjects used the corresponding hand to make choices. Note that subjects learned the definition of high- and low-pitch categories in a pretest, i.e., listening to 180 Hz and 360 Hz pure tones.

**Figure 1.**
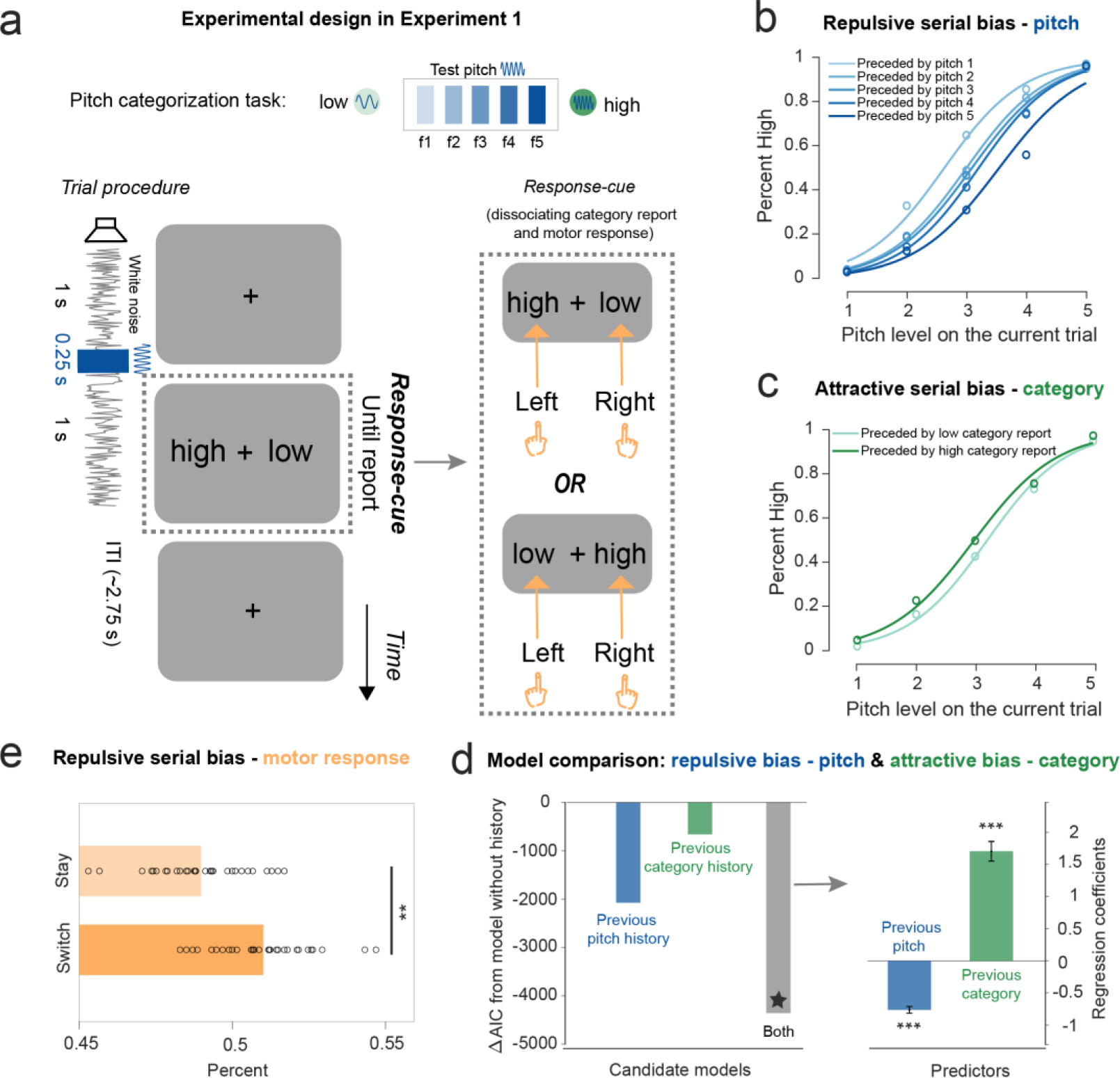
Task paradigm and behavioral serial bias in Experiment 1. **A**. Auditory categorization task paradigm. Upper: pitch of the pure tone stimulus was pseudo- randomly selected from 5 fixed frequencies (f1, f2, f3, f4, f5) between 180 Hz (low category; light green) and 360 Hz (high category; dark green). Lower: In each trial, subjects categorized a given 0.25 s pure tone (dark blue; selected from f1, f2, f3, f4, f5) embedded in a 2.5 s sustained white noise stream (grey line) into “high” (closer to 360 Hz) or “low” (closer to 180 Hz) category. After the pure tone, a response-cue screen appeared, based on which subjects used the corresponding hand to made responses (right inset). **B**. Pitch serial bias (aggregate results across subjects). “High” category report percent as a function of the current pitch, for different pitch in previous trial (Light- to-ark color lines denote low-to-high pitches). Each circle represents aggregated data for each condition. Solid lines represent the logistic regression fits. **C**. Category serial bias (aggregate results across subjects). “High” category report percent as a function of the current pitch, for different category report in the previous trial (light green: low category report; dark green: high category report). **D**. Model comparison results. Left: ΔAIC of Model 2 (blue; current trial + previous pitch), Model 3 (green; current trial + previous category report), and Model 4 (grey; current trial + previous pitch + previous category), compared to Model 1 (current trial only). Right: Regression coefficients for previous pitch (blue) and previous category report (green) extracted from the winning model (Model 4, * in model comparison). Error bars represent 95% confidence interval. **E**. Motor response serial bias (left or right hand). “Switch” trials (dark orange): percentage of trials that are different from previous trial in motor response. “Stay” trials (light orange): percentage of trials that are the same as previous trial in motor response. Each circle represents individual subject. (**: p < 0.01, ***: p < 0.001).

As shown in Fig. 1BC, both pitch and category exhibited the serial dependence effect, yet in different directions. Specifically, the pitch of the current tone tends to be perceived away from that of the preceding pitch, i.e., repulsive serial bias (Fig. 1B). For example, the same f3 on the current trial was more likely categorized into the “high” category when preceded by lower pitch (light blue) compared to when preceded by higher pitch (dark blue). In contrast, the categorization performance displayed an attractive serial bias, such that the same f3 on the current trial was more likely to be reported as the “high” category when the previous report was also “high” (Fig. 1C).

Four generalized linear mixed-effects models (GLMMs) were built to account for the behavioral performances. Model 1 assumes that the category report only depends on the current pitch and a constant bias (base model; no serial bias). Model 2 and 3 consider additional contributions from previous pitch or previous category report, respectively. Model 4 takes account of contributions from both pitch and category reports in the preceding trial. Model 2 – 4 were evaluated against Model 1 by comparing the Akaike information criterion (AIC) values. As shown in Fig. 1D (Left), all the three models were better than Model 1 ((Model 2: ΔAIC = -2079; Model 3: ΔAIC = -657; Model 4: ΔAIC = -4369), indicating the influence of past-trial information on current perception. Moreover, Model 4 outperformed Model 2 and Model 3, supporting that pitch and category history together affect the current category decision.

Importantly, as shown in Figure 1D (right panel), Model 4 showed negative coefficients for past-trial pitch (mean = -0.79; 95% CI = [-0.89, -0.69]; t(58002) = -15.44, p < 0.001, one-sample t-test), but positive coefficients for past-trial category report (mean = 1.70; 95% CI = [1.41, 1.99]; t(58002) = 11.59, p < 0.001, one-sample t-test), confirming the repulsive and attractive serial bias for pitch and category report, respectively. Furthermore, the motor response, which is designed to be independent of category report, also exhibited serial bias, in a repulsive manner (Fig. 1E). Specifically, the “switch” motor response probability across consecutive trials was greater than non-switching (“Stay”) (“Switch”: mean = 0.51, SD = 0.016; “Stay”: mean = 0.49, SD = 0.016; “Switch” vs. “Stay”: paired-sample t-test, t(29) = 3.49, p = 0.0015).

Together, subjects’ perceptual decision tends to be concurrently shifted by multiple features on the preceding trial, being repulsed from the previous pitch while attracted to prior category reports.

### Decoding multiple features of the current trial

We first performed a time-resolved multivariate decoding analysis^18, 25–27^ to examine the neural representations of multiple features of the current trial – Pitch, Category, Motor response. A linear regression analysis was used to fit the neural dissimilarity to feature dissimilarity, and the time-resolved regression coefficients denote the decoding performance, for each feature, at each time point, and in each subject (see details in Methods).

As shown in Figure 2A, the pitch (blue) and reported category (green) of the current trial could be decoded shortly after the pure tone (cluster-based permutation test, p < 0.001, one-sided, corrected; Pitch: 84-1144 ms after tone onset; Category: 84 to 934 ms after tone onset). Furthermore, the motor response decoding (Fig. 2A, orange) occurred after the response cue frame (cluster-based permutation test, p < 0.001, one-sided, corrected; Motor response: 44-994 ms after response cue frame). This is well expected since subjects could only determine the responding hand after the response cue.

**Figure 2.**
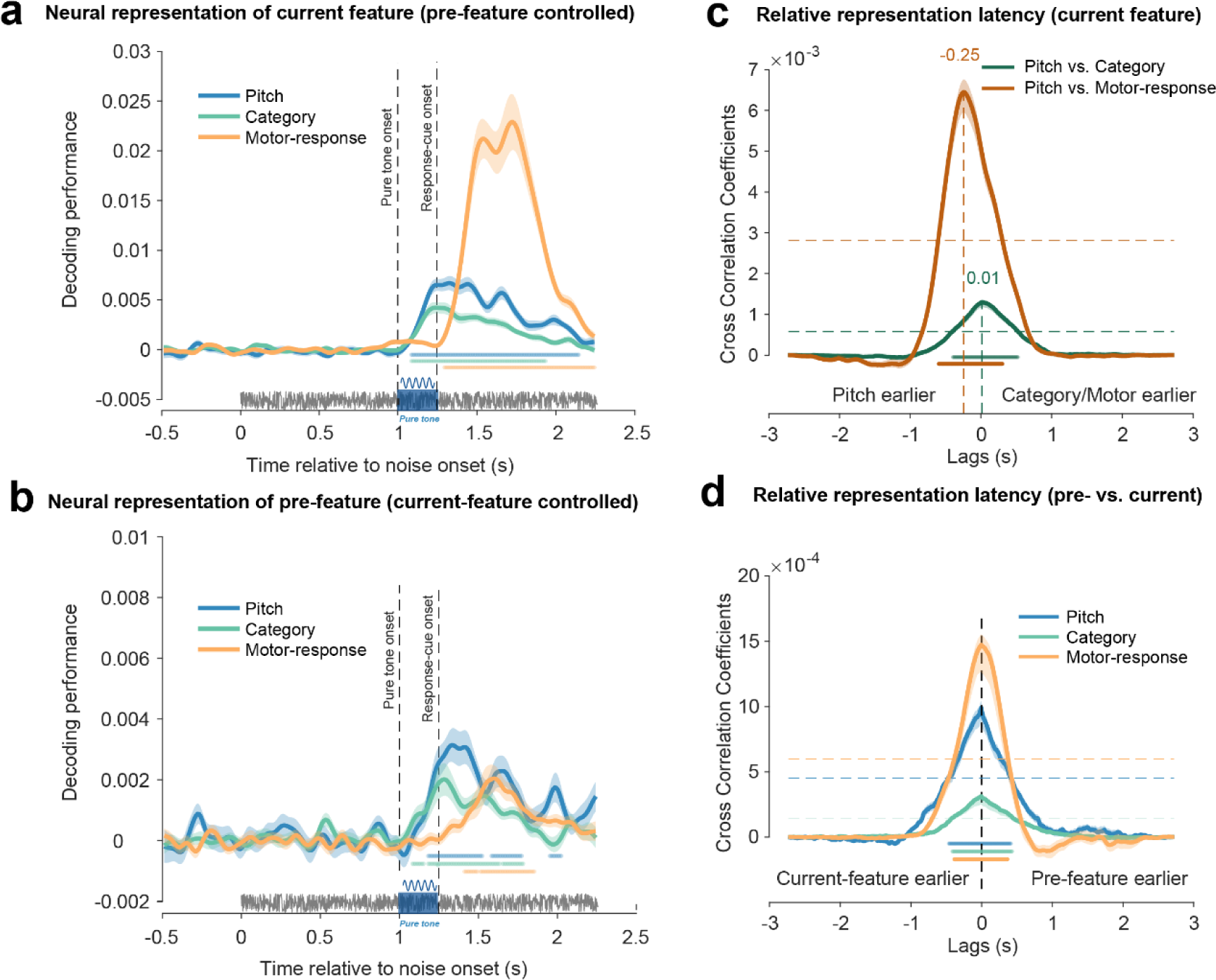
Co-occurrence of past-trial neural reactivations and neural encoding of current-trial features. **A.** Grand average decoding performance for current-trial features as a function of time following the sustained white noise, for Pitch (blue), Category report (green), and Motor response (orange). The pure tone (blue rectangle) was embedded in a 2.25 sec sustained white noise (grey horizontal line). Vertical dashed lines from left to right denote the tone onset and response cue frame, respectively. Horizontal colored lines denote significant temporal clusters (cluster-based permutation test, p < 0.001, one-sided, corrected) for each feature. Shadows represent SEM. **B.** The same as A but for past-trial features. Note that the past-trial decoding is performed by fixing the current-trial feature. **C.** Cross-correlation coefficients as a function of temporal lag for each current-trial feature pair (Green: Pitch vs. Category; Brown: Pitch vs. Motor response). Shadow represents 95% confidence interval ((bootstrapping, N = 5000). Horizontal colored lines denote significant temporal clusters (cluster-based permutation test, one-sided, corrected, p < 0.05) for each feature pair. Horizontal dashed lines represent the 95% permutation threshold (corrected for multiple comparison). Vertical dashed line indicates the significant peak indexing the relative activation lag. **D.** Cross-correlation coefficients between past-trial reactivation and current-trial neural response, as a function of temporal lag, for each feature (Blue: Pitch; Green: Category report; Orange: Motor response). The peak at 0 ms temporal lag (vertical dashed line) supports the co-occurrence of past-trial reactivation and current-trial neural response.

Overall, multiple features of the current trial could be successfully decoded from the neural response, i.e., the pitch and category information about the pure tone emerges right after the tone onset, and the neural code of motor response arises following the response cue frame.

### Past-trial features are reactivated by corresponding events in the current trial

Interestingly, as shown in Figure 2B, past-trial features, i.e., pitch (blue), category report (green), and motor response (orange) could also be decoded form the neural response of the current trial. Crucially, to exclude potential current-trial confounding when decoding past-trial features, we performed the past-trial decoding analysis for each of the same current features and then combined the results (see Methods for details).

Most importantly, we found that past-trial features were reactivated by the corresponding event in the current trial (Fig. 2B). Specifically, previous-pitch decoding (blue line, Fig. 2B) was at chance level prior to the pure tone and rose right after tone onset (cluster-based permutation test, one-sided, corrected; significant clusters: 184-524 ms, 584-774 ms, and 954-1024 ms, p < 0.05). Similarly, previous-category decoding (green line, Fig. 2B) emerged after the tone onset (cluster-based permutation test, one-sided, corrected, significant clusters: 84-154 ms, 184-634 ms, and 654-784 ms, p < 0.05). In contrast, rather than being triggered by the pure tone, previous-motor-response (orange line, Fig. 2B) occurred after the response cue frame (cluster-based permutation test, one-sided, corrected; clusters: 164-244 ms and 264-604 ms after the response cue onset, p < 0.05).

Overall, features of the preceding trial occurring seconds before and retained in WM are reactivated by specific events in the current trial, i.e., pitch and category by the tone stimulus and motor response by the response cue. It is noteworthy that past-trial features, given their maintenance in WM, could potentially be reactivated by any triggering event in the current trial, yet the findings support a feature-specific reactivation temporal profile.

### Co-occurrence of past-trial reactivations and neural coding of current-trial features

We next examined the temporal relationship between the current-trial neural responses and past-trial reactivations, by calculating their cross-correlation coefficients over time. First, features in the current trial showed varied latencies in their neural response (Fig. 2C). Specifically, the Pitch vs. Category (green line) correlation coefficient was significant from -400 to 520 ms time lag (permutation test, one-sided, corrected, p < 0.05), peaking at 10 ms lag (bootstrapping, 95% CI = [0, 70] ms, N = 5000), while the Pitch vs. Motor response (Brown line) showed peak around -250 ms lag (bootstrapping, N = 5000, 95% CI = [-270, -230] ms; permutation test, one-sided, corrected, p < 0.05, - 600 to 300 ms). Thus, pitch lagged category by 10 ms and led motor response by 250 ms. The latter result is well expected given the 250 ms interval between pure tone and response cue frame.

Most importantly, as shown in Figure 2D, all the three features showed temporally aligned profiles between current-trial neural responses and past-trial reactivations (permutation test, one-sided, corrected, p < 0.05; Pitch: blue, -460 to 420 ms; Category: green, -410 to 440 ms; Motor response: orange, -380 to 370 ms), peaking at 0 ms time lag (bootstrapping, N = 5000, 95% CIs = [0, 0] ms for Pitch and Category, and [-60, 110] ms for Motor response), advocating simultaneous activation of present and past information for each feature in the present trial.

Thus, the reactivation of past-trial information occurs concurrently with the activities of present information. In other words, each feature’s current and past information arise simultaneously in the present trial, which potentially leads to the past-to-present influence and engenders serial bias.

### Past-trial feature shifts neural representation of current-trial feature modulating serial bias behavior

After revealing the co-emergence of past-trial reactivations and current-trial neural responses (Fig. 2), we next examined the direct neural evidence for serial bias. Specifically, we developed a novel analysis by accessing whether the neural representation of the current-trial feature would be attracted toward or repulsed from the preceding feature.

Figure 3A (left panel) exemplifies the general idea to test pitch serial bias, e.g., quantifying the influence of past-trial f2 on current-trial f1. First, we built the neural templates for f1 (dark blue) and f2 (light blue) based on all trials.

**Figure 3.**
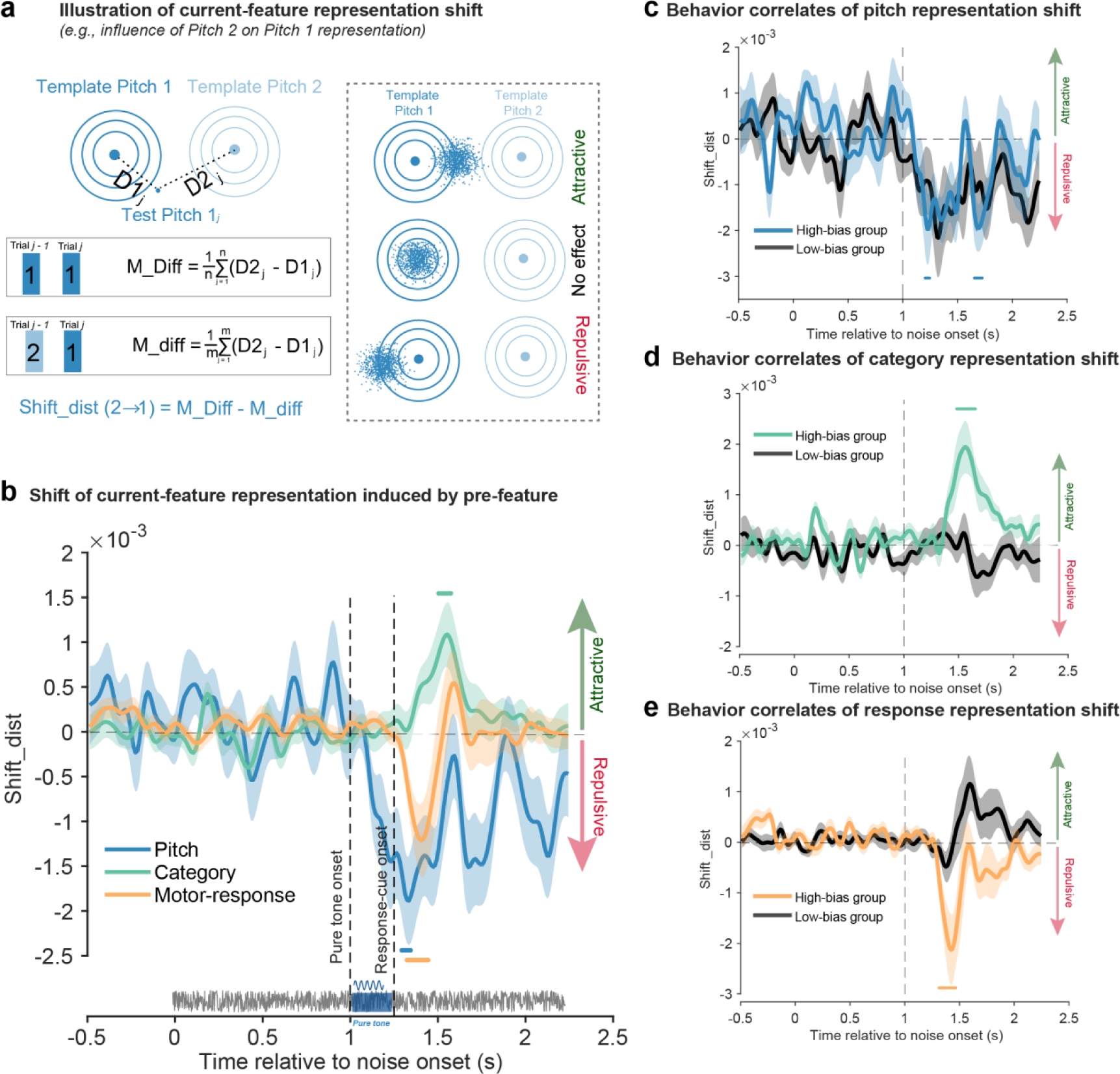
Direct neural evidence for serial bias and its behavioral relevance. **A.** Illustration of neural representational shift analysis. Left: influence of previous f2 on current f1 as an example. Neural templates were built for f1 (dark blue circle) and f2 (light blue circle) base on all trials. For each trial with current f1 and previous f2, its neural distance to template f1 and template f2 were computed, resulting in D1, D2, and their difference M_diff (D2-D1). As baselines, trials that have current f1 but previous f1 were selected and were analyzed using the same analysis, resulting in M_Diff. The difference between M_Diff and M_diff (Shift_dist) characterizes the neural representational shift in f1 by prior f2. Right: Neural representation of current-trial f1 is attracted toward (upper, positive Shift_dist values), repulsed from (lower, negative Shift_dist values), or not affected (middle, around zero Shift_dist values) by prior f2. **B.** Grand average neural representational shift (Shift_dist) as a function of time following white noise onset, for Pitch (blue), Category report (green), and Motor response (orange). Positive and negative values correspond to attractive and repulsive direction, respectively. Horizontal colored lines denote significant temporal clusters (cluster-based permutation test, two- sided, corrected, p < 0.05) for each feature. Shadows represent SEM. **C-E.** Subjects were divided into two groups based on the serial bias behavior in Pitch, Category, or Motor response, respectively. Grand average neural representational shift (Shift_dist) of High-bias (colored lines) and Low-bias (black line) groups for Pitch (C), Category (D), and Motor response (E). Horizontal colored lines denote significant temporal clusters (cluster- based permutation test, one-sided, corrected, p < 0.05).

Next, we chose trials (current f1, previous f2) and computed the neural distance between each trial in the subsample and the f1 and f2 templates, separately, yielding D1 and D2, from which M_diff (D2-D1) was obtained. For comparison, we chose trials (current f1, previous f1) as baselines and did the same distance computation, yielding the M_Diff (D2-D1). Finally, the difference between M_diff and M_Diff was calculated (Shift_dist) to quantify the influence of the previous f2 on the neural representation of current f1. If previous f2 attracts f1 neural representation, we would expect smaller M_diff than M_Diff, yielding positive Shift_dist values, and vice versa. There are three possibilities – attractive, no effect, and repulsive (Fig. 3A, right panel), corresponding to positive, around zero, and negative Shift_dist values, respectively. A similar idea has been applied to category and motor response (see Methods for details). The neural shift analysis was performed at each time point, for each feature, and in each subject.

Figure 3B plots the neural shift for the three features, with positive and negative values denoting attraction towards and repulsion from the past feature, separately. Pitch (blue line) showed a repulsive bias, rising shortly after pure tone (cluster-based permutation test, two-sided, corrected; 294-344 ms after the tone onset, p = 0.0056), consistent with the repulsive direction in serial bias behavior (Fig. 1BD). In contrast, the category (green line) displayed an attractive shift towards previous information, occurring relatively late (cluster-based permutation test, two-sided, corrected; 504-574 ms after the tone onset, p = 0.0058), congruent with the attractive bias in behavior (Fig. 1CD). Finally, the neural code of motor response (orange line) was shifted away from that of the prior trial (cluster-based permutation test, two-sided, corrected; 74-194 ms after the response cue, p = 0.005), again in line with behavior (Fig. 1E). Thus, the findings demonstrate direct neural evidence for serial bias, revealing a behaviorally congruent shift in neural representation for pitch, category, and motor response.

Finally, we assessed the behavioral relevance of the neural shift. For each feature, all the participants (N = 30) were divided into two groups of the same size – High-bias and Low-bias– based on their serial bias in behavior (see Methods for details). The High-bias group displayed a significant neural shift for pitch (Fig. 3C, blue; cluster-based permutation test, one-sided, corrected, 204-244 ms and 654-724 ms after the tone onset, p < 0.05), category (Fig. 3D, green; 484-664 ms after the tone onset, p < 0.001), and motor response (Fig. 3E, orange; 64-224 ms after the response cue onset, p < 0.001), while not for the Low-bias group (Fig., 3CDE, black line).

Taken together, we demonstrate direct neural evidence for serial bias, revealing a behaviorally congruent shift in neural representation by past-trial information, in a feature-specific way, and the neural shifting profiles further correlate with serial bias behavior.

### Past-trial reactivations are not due to early task-relevant events or temporal prediction (Experiment 2)

In Experiment 1, pitch and category information of the previous trial was reactivated by the corresponding event, i.e., pure tone (Fig. 2B). Meanwhile, since the pure tone occurred early in each trial (Fig. 1A), it might be the first task-relevant event (i.e., pure tone) rather than the feature-specific event that reactivated the past-trial information. To test the possibility and also confirm the findings of Experiment 1, we designed Experiment 2 (N = 30) during which subjects performed the same auditory categorization task as Experiment 1, except now the response-cue frame appeared at the beginning of each trial (Fig. 4A). If the pith and category reactivations are indeed due to the early task-relevant event, we would expect their reactivations right after the response-cue frame. Moreover, since the white noise occurred at a fixed temporal lag after the response-cue frame in Experiment 2 (Fig. 4A), we could also test whether the reactivation simply derives from temporal prediction.

**Figure 4.**
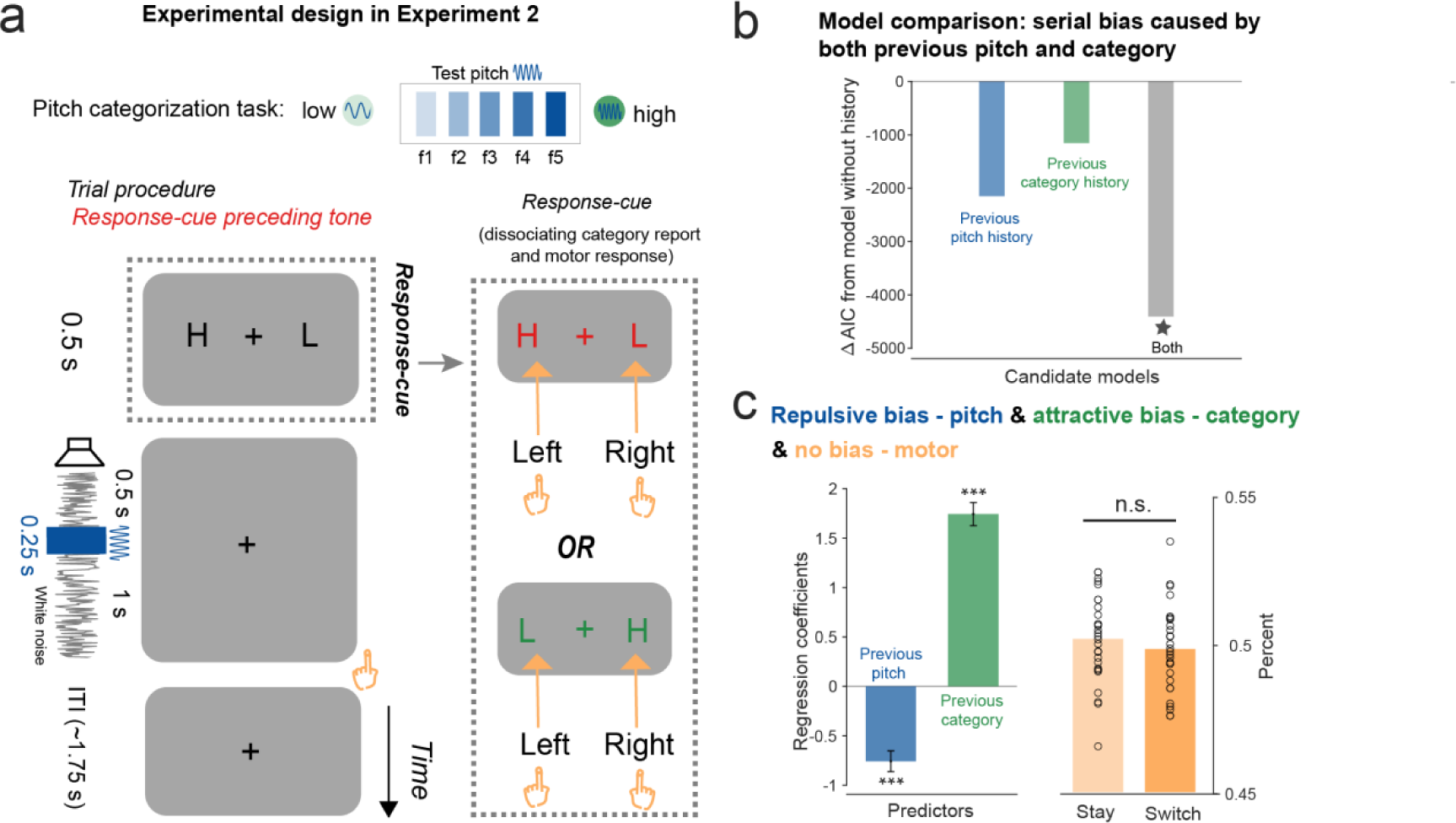
Task paradigm and behavioral serial bias in Experiment 2. **A**. Experiment 2 employed the same auditory categorization paradigm as Experiment 1, with only one difference, i.e., the response-cue screen appeared before the white noise (dotted box). Subjects categorized the following 0.25 s pure tone (dark blue) embedded in a 1.75 s sustained white noise (grey line) into “high” or “low” category, by using the corresponding hand defined in the response-cue frame (dotted box) appeared at the beginning of each trial. **B**. Model comparison results. ΔAIC of Model 2 (blue; current trial + previous pitch), Model 3 (green; current trial + previous category report), and Model 4 (grey; current trial + previous pitch + previous category), compared to Model 1 (current trial only). **C.** Left: Regression coefficients for previous pitch (blue) and previous category report (green) extracted from the winning model (Model 4, * in model comparison). Error bars represent 95% confidence interval. Right: No motor response serial bias, i.e., no difference between “Switch” trials (dark orange) and “Stay” trials (light orange). Each circle represents individual subject. (***: p < 0.001).

First, subjects exhibited similar serial bias behavior as Experiment 1. All the history-dependence models outperformed the history-free model (Fig. 4B; Model 2 vs. Model 1: ΔAIC = -2148; Model 3 vs. Model 1: ΔAIC = -1151; Model 4 vs. Model 1: ΔAIC = -4406). Furthermore, the pitch and category report showed negative (regression coefficient: mean = -0.76; 95% CI = [- 0.96, -0.55]; t(58196) = -7.24, p < 0.001, one-sample t-test) and positive serial bias (mean = 1.74; 95% CI = [1.51, 1.97]; t(58196) = 14.90, p < 0.001, one-sample t-test), respectively (Fig. 4C, left). Meanwhile, Experiment 2 did not show serial bias for motor response (Fig. 4C right, “Switch”: mean = 0.50, SD = 0.014; “Stay”: mean = 0.50, SD = 0.014; “Switch” vs. “Stay”: paired-sample t-test, t(29) = -0.66, p = 0.52), consistent with previous results^23^.

Importantly, as shown in Figure 5, Experiment 2 did not show an early reactivation of past-trial information after the response-cue frame or the white noise, but instead largely replicated Experiment 1. First, neural representation of the current-trial features emerged after the tone stimulus (Fig. 5A, cluster-based permutation test, p < 0.001, one-sided, corrected; -26-834 ms for Pitch; 14 to 884 ms for Category; 44-1044 ms for Motor response), with similar latencies (Fig. 5B; Pitch vs. Category: correlation coefficients, 0 ms lag, bootstrapping, 95% CI = [-10, 20] ms, N = 5000; Pitch vs. Category: permutation test, one-sided, corrected, -360 to 400 ms; Pitch vs. Motor: correlation coefficients, -40 ms lag, bootstrapping, N = 5000; Pitch vs. Motor: 95% CI = [-150, 0] ms, permutation test, one-sided, corrected, p < 0.05, - 490 to 360 ms). Second, past-trial pitch (blue) and category (orange) features could also be decoded from neural response (Fig. 5C; cluster-based permutation test, one-sided, corrected; Pitch: 154-554 ms and 574-674 ms after tone onset, p < 0.05; Category: 84-584 ms and 714-824 ms after tone onset, p < 0.040). Critically, similar to the findings of Experiment 1, past-trial reactivations co-occurred with the current-trial neural response of features, for both pitch and category (Fig. 5D; cross-correlation coefficients: 0 ms lag for both Pitch and Category, bootstrapping, 95% CIs = [0, 0] ms, N = 5000; permutation test, one-sided, corrected, Pitch: -390 to 380 ms, Category: -340 ms to 400 ms).

**Figure 5.**
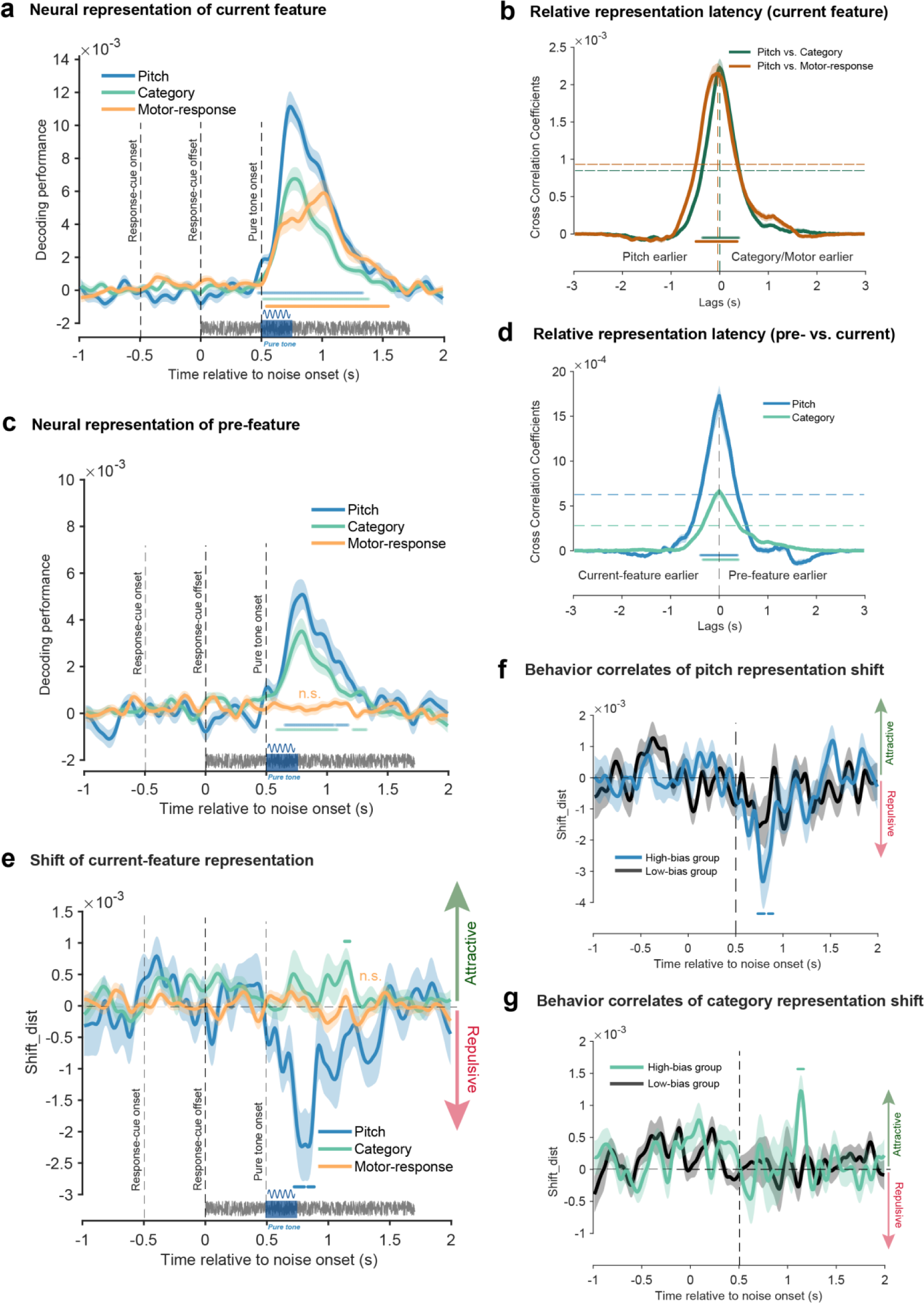
Neural results of Experiment 2. **A.** Grand average decoding performance for current-trial features as a function of time following the sustained white noise, for Pitch (blue), Category report (green), and Motor response (orange). Note that the response- cue frame occurred before white noise onset (-0.5 s, dashed vertical line). Time 0 denotes the white noise onset. Vertical dashed lines from left to right denote the onset and offset of response-cue frame, and the pure tone onset, respectively. Horizontal colored lines denote significant temporal clusters (cluster-based permutation test, p < 0.05, one-sided, corrected) for each feature. Shadows represent SEM. **B.** Cross- correlation coefficient time course as a function of temporal lag, for each current-trial feature-pair (Green: Pitch vs. Category; Brown: Pitch vs. Motor response). Shadow represents 95% confidence interval (bootstrapping, N = 5000). Horizontal colored lines denote significant temporal clusters (permutation test, one-sided, corrected, p < 0.05) for each feature pair. Horizontal dashed lines represent the 95% permutation threshold (corrected for multiple comparison). Vertical dashed line indicates the peak correlation coefficient indexing the relative activation lag. **C.** The same as A but for past-trial features. **D.** Cross-correlation coefficients between past-trial reactivation and current-trial neural response, as a function of temporal lag, for Pitch (Blue) and Category (Green).

Finally, Experiment 2 also replicated the behaviorally-relevant neural shifting results, for both pitch and category (Fig. 5E). Specifically, neural representations of the current-trial pitch (blue) and category (green) displayed repulsive and attractive shifts by previous features, respectively (cluster-based permutation test, one-sided, corrected; Pitch: 224-314 ms and 334-384 ms after the tone onset, p < 0.01; Category: 634-674 ms after the tone onset, p = 0.056). Moreover, the neural shifts were correlated with serial bias behavior for both pitch (Fig. 5F) and category (Fig. 5G), i.e., showing significant neural shifts for the High-bias but not Low-bias group (cluster-based permutation test, one-sided, corrected; Pitch: 234-304 ms and 344-394 ms after tone onset, p < 0.05; Category: 594-674 ms after tone onset, p = 0.017). Notably, consistent with the lack of behavioral serial bias for motor response (Fig. 4C), we found neither significant past-trial reactivation (Fig. 5C, orange) nor neural shift for motor response (orange, Fig. 5E).

Together, past-trial features could not be reactivated by either an early task-relevant event (i.e., response-cue frame) or a temporally predictable stimulus (i.e., white noise). Instead, only the corresponding feature-specific event could trigger the past-trial reactivations, with their co-occurrences giving rise to serial bias.

The peak at 0 ms temporal lag (vertical dashed line) supports their co-occurrence. Other denotations are the same as B. **E.** Grand average neural representational shift (Shift_dist) as a function of time relative to the white noise onset, for Pitch (blue), Category report (green), and Motor response (orange). Positive and negative values correspond to attractive and repulsive direction, respectively. Horizontal colored lines denote significant temporal clusters (cluster-based permutation test, one-sided, corrected, p < 0.05 for Pitch, p = 0.056 for Category) for each feature. Shadows represent SEM. **F-G.** Subjects were divided into two groups based on the serial bias behavior in Pitch and Category, respectively. Grand average Shift_dist of High-bias (colored lines) and Low-bias (black line) groups for Pitch (F) and Category (G). Horizontal colored lines denote significant temporal clusters (cluster-based permutation test, one-sided, corrected, p < 0.05).

## Discussion

In two auditory categorization experiments, we reveal a new “event-file” reactivation neural mechanism underlying serial bias that occurs spontaneously across trials. Specifically, we demonstrate that serial bias arises from the co-emergence of past reactivation and the neural coding of present inputs, which renders their interactions and the shifting of current information. Importantly, we provide converging behavioral and new neural evidence that the serial bias occurs in a feature-dissociated manner, i.e., lower-level (i.e., pitch and motor response) and higher-level (i.e., category) features are repulsed from and attracted to the corresponding ones in the previous trial, respectively, within the same task. Moreover, the past-trial reactivation is not due to either early task-relevant events or temporal prediction but triggered by the occurrence of the feature-specific event. Taken together, features in the current “event-file” automatically trigger the corresponding WM traces of past trials, and their co-occurrence leads to serial bias. This “event-file” reactivation might reflect a fundamental operational mechanism for past-to-present adaptive optimization.

The past-to-present influences occur in a wide range of paradigms and contexts ^2, 28, 29^, of which the serial bias represents an extreme phenomenon. It refers to the involuntary, systematic shifting of current perception by previous trials and appears on many features ^3–11, 30–">33^. The typical serial dependence effect is in an attractive direction^3, 11, 30–32^ while the repulsive bias and individual difference have also been observed ^7, 8, 10, 11, 34, 35^. The incongruency might be due to the high correlations of features in previous experimental designs; as a result, the overall effect would be a mixture of feature-specific serial biases with varied directions^5, 7, 33^. Here by carefully disentangling features in the design, we revealed, within the same task, the dissociated serial bias for pitch, category, and motor response, thus reconciling previous contradictions.

Furthermore, we found that pitch and motor response were repulsed from past information, while the category displayed an attractive direction, also in line with the view that repulsive and attractive serial biases are associated with perceptual and post-perceptual processing levels, respectively^5, 8, 36–38^. Most importantly, we provided direct neural evidence for serial bias, i.e., the behaviorally congruent and relevant shifting of neural representation toward or away from the past at different latencies, and further disclosed a new neural mechanism underlying serial bias.

For the past to influence the present, either voluntarily or spontaneously, prior information should leave traces in WM. It has long been viewed that the WM process relies on persistent firing^39–42^ or frequency-specific neural oscillations^43, 44^. Interestingly, information could also be retained within synaptic weights of WM networks via short-term neural plasticity principles and thus in an ‘activity-silent’ way ^12–17^. However, recent studies suggest that the ‘activity-silent’ mode could only passively maintain information^13^ (including ours, see ref^45–47)^, and memory manipulation still relies on the reactivations of WM to active states^19, 20^. As a consequence, for serial bias to occur, the past WM traces, retained either actively or silently, should encounter the current inputs in the time to alter the present. Consistent with the view, we demonstrate the co-emergence of past reactivation and present information, followed by the shifted neural representation of current features. Moreover, the reactivation profiles encompass multiple feature-specific reactivations that arise simultaneously as the corresponding features in the current trial.

Notably, past-trial reactivation is not due to temporal expectancy. In Experiment 2 when the white noise appeared at a fixed time after the response-cue frame in each trial, the pitch and category reactivations were still triggered by the auditory tone rather than the white noise. Together, the “event-file” WM traces are reactivated by corresponding features in the current trial so that the feature-specific co-occurrence in neural space contributes to the serial bias.

A recent interesting study revealed brief reactivation of past spatial information during the inter-stimulus interval which was further correlated with serial bias behavior^21^. Here we observed the concurrent emergence of past reactivation and present information after the stimulus. Both findings thus support the reactivation of past features from activity-silent to activity-based WM state for serial bias to occur, yet within different temporal ranges, i.e., pre-stimulus baseline vs. after-stimulus. Latent memory traces have been suggested to arise from the silent state whenever needed ^48, 49^. Accordingly, the prominent pre-stimulus reactivation shown in the previous study might be due to the strong temporal anticipation given the fixed inter-trial interval (ITI), while our study randomized the ITI across trials and therefore did not observe strong pre-stimulus signatures. Moreover, our alpha-band decoding control analysis showed that information about past category and motor response (but not for pitch) was indeed present before tone onset (supplementary Fig. 3), thus partially in agreement with previous results. Moreover, as information might be carried via neural oscillations during the activity-silent state ^23, 24, 43, 44^, the past reactivation might also reflect an enhancement of weakly coded memory traces via decreasing cross-trial variability^50^. Nevertheless, reactivation by themselves, either before or after stimulus, could not fully explain the serial bias, and here we provide direct neural evidence that current information is indeed shifted towards or away from past (i.e., attractive, repulsive) following their co-occurrence.

Importantly, we extend previous single-feature emphasis to the “event-file” framework, within which multiple features and their respective serial biases are disentangled in both behavioral performance and neural representations. It is recognized that even the simplest perceptual decision task encloses packs of features, e.g., physical properties, abstract categories, response actions, etc., collectively constituting an “event-file” ^22^. Here we show that within the same auditory categorization task, the past “event-file” is implicitly imprinted in WM and automatically passed to the next “event-trial” in a feature-encapsulated way. Importantly, not any events can trigger past information, also in line with previous findings ^46, 47^. For instance, white noise, despite being an auditory sound within the same sensory modality, failed to reactivate prior pitch and category information, and previous motor response, although apparently retained in WM, could not be triggered by either white noise or tone stimulus. Thus, past trial automatically leaves a memory trace of “event-file” within which multiple features keep their identities and specificities, exerting their respective impacts on the future.

Serial bias denotes a dramatic case about past-to-present influence since information in the previous trial, which occurs several seconds ago and in principle should be discarded, still biases the present trial. The phenomenon is therefore independent of several factors involved in many other paradigms, such as voluntary attention, task-relevant modulation, task-irrelevant capture, etc^13, 48, 51^. Moreover, as trials are typically several seconds apart with random ITI in-between, serial bias could not arise from within-trial temporal effects either, such as priming, adaptation, etc ^28, 29^. Furthermore, while potentially sharing similar WM storage underpinnings, serial bias essentially differs from WM studies that would instruct subjects to at least retain certain information. Our findings thus implicate a presumably ubiquitous mechanism for temporal dependence in many cognitive processes, such as perception, attention, memory, and decision making.

Taken together, every single present is intertwined with the past, yet the past-to-present influence is not feature-agnostic but feature-specific, allowing for information encapsulation through trial-by-trial updates. The current feature reactivates the corresponding traces left in WM, and their co-occurrence induces the neural interactions and generate serial bias, thereby facilitating automatic adaptive generalizations from past to present.

## Methods

### Experiment procedures and subjects

#### Participants

Thirty subjects (20.8 ± 2.2 years old, 12 females) took part in Experiment 1, and thirty new subjects (20.6 ± 1.7 years old, 14 females) participated in Experiment 2. The sample size was determined based on previous serial dependence behavioral studies^7^. They are all right-handed with normal or corrected to normal vision and normal audition. Participants were paid and gave written informed consent before starting the experiment. The study was conducted in accordance with the Declaration of Helsinki, and was approved by the Ethical Committee of the School of Psychological and Cognitive Sciences at Peking University.

#### Apparatus

Experimental programs were developed using Matlab (MathWorks Inc., Natick, MA, USA) and Psychophysics Toolbox^52^. Auditory stimuli were controlled by external sound card RME Babyface pro and were emitted through Sennheiser CX213 earphones. The loudness of sound was adjusted to a comfortable level (∼65 dB SPL). The visual stimuli were presented on a Display++ LCD screen with a refreshing rate of 120 Hz and a resolution of 1280 × 1024 pixels. Participants sat at a distance of 100 cm from the screen and their heads were maintained steady using a chin-rest. In the main task, a Cedrus RB-540 response box was used to collect participants’ responses.

#### Stimuli and experimental procedure

The experiment consisted of two phases, a behavioral training phase, and a formal testing phase. In the behavioral training phase, participants had to memorize two tones, a “low” tone (180 Hz) and a “high” tone (360 Hz), without EEG recordings, so that the “Low-pitch” and “high-pitch” categories could be successfully formed before the formal test. In the formal testing phase, participants performed a categorization task, reporting whether a given auditory tone is more similar to the memorized “low” tone or “high” tone, i.e., an auditory categorization task, with EEG recordings.

#### Behavioral Training phase

We designed two types of tasks to help participants memorize these two tones (see supplemental Figure S1): a reproduction task and a recall task. In the reproduction task, on each trial, the pure tone (250 ms, 10 ms ramp-up, ramp-down time) was presented first and participants were required to reproduce it afterward. In the recall task, a recall cue (“low” or “high”) was presented to participants at the beginning of each trial. They were asked to produce it according to their memory. The training phase consisted of two sessions. The first session is a block-wise design with four ten-trial blocks containing a “high” tone reproduction task, “high” tone recall task, “low” tone reproduction task, and “low” tone recall task respectively. The order of “high” or “low” tones is counterbalanced among participants. In the second session, the stimuli were trial-by-trial randomized. This session had two forty-trial blocks, with the first block containing a reproduction task with feedback and the second block containing a recall task without feedback.

For the reproduction task, participants were presented with a 250 ms pure tone (180 Hz or 360 Hz), and 3 seconds later, they adjusted the pitch of a probe tone which was randomly selected from 13 initial frequencies (mean is the probe frequency, log step 0.1) with up and down arrows (log step 0.1) to match the frequency to the target tone. For the recall task, a retro-cue (“high” or “low”) was presented for 2 seconds. Participants then used up and down arrows to adjust the tone frequency to match the target tone (180 Hz or 360 Hz). The inter-trial interval was 1 to 1.5 s. Participants moved to the testing phase only when they accomplished the training phase tasks.

Participants with poor recalling performance in the training session would be excluded from participating in the formal test.

#### Formal testing phase

Participants had to compare the probe tone to the memorized “low” tone (180 Hz) and “high” tone (360 Hz) and chose whether the tone was more similar to the “low” or “high” tone. To equalize the task difficulty across participants, we individualized the probe tones. First, we used the same five probe tones (202, 227, 255, 286, and 321 Hz) to test each participant’s performance. Then we fitted a logistic regression to the percentage of “high” reports as a function of pitches and acquired pitches (f1, f2, f3, f4, f5) corresponding to percents 0.1, 0.3, 0.5, 0.7, and 0.9 for each subject. Then we employed these five pitches in the main task when EEG signals were recorded.

***Experiment 1*** In each trial, the probe tone embedded in white noise was presented while participants kept their eyes on the central cross. The white noise lasted 2.25 s and the probe tone (duration, 0.25 s) was presented 1 s after noise onset. After the offset of the probe tone, a response cue was immediately presented on the screen, based on which participants chose the corresponding hand to make a response. For example, if the “high” and “low” characters were presented on the left and right side respectively, participants had to press a left button for “high” and a right button for “low” categorical choice, and vice versa. Participants were required to respond as accurately and fast as possible. The inter-trial interval is 1.5 s to 2 s. The formal test consisted 20 blocks with 100 trials for each block. Five tones (f1, f2, f3, f4, f5) were pseudo-randomized within each block (20 trials for each pitch).

Participants performed 2000 trials in total, which were divided into two sessions accomplished on two different days.

***Experiment 2*** The stimuli and task were similar to those in Experiment 1, except that the response cue was presented before the tone onset, and that the white noise lasted 1.75 s and the probe tone (duration, 0.25 s) appeared 0.5 s after noise onset. Specifically, at the beginning of each trial, a response cue was presented for 0.5 s, which was followed by the auditory stimuli. Participants chose the corresponding hand to make a response based on the response cue. The response cue was denoted with letters on screen and colors (red or green).

### Behavioral data analysis and modeling

#### Aggregate analysis

To examine the pitch serial bias, i.e., how current categorical decisions are influenced by pitches in the previous trial, we first combined data from all participants into aggregated data. The aggregated data were then divided into five groups based on the previous pitch, and then for each group, we calculated the percentage of “high” reports for each pitch on the current trial. Similarly, to examine the serial bias for category reports, i.e., how current categorical decisions are influenced by category reports on the previous trial, data were divided into two groups based on the previous category report.

Finally, we also examined whether there is a serial bias in motor response by comparing the switching rate (percentage of trials with motor responses different from the previous response) and staying rate (1-switching rate).

#### GLM models

To visualize the overall pattern of serial bias, for each group of the aggregate data, we first fitted a logistic regression model (glmfit function in the Matlab Statistics and Machine Learning Toolbox) to characterize the relationship between the percentage of “high” reports and the pitch on the current trial.

To quantitatively test the serial bias caused by previous pitch and its corresponding category, we built four generalized linear mixed-effects models (GLMMs) with model 1 assuming no history influence, models 2 and 3 assuming previous pitch and previous category affect current decisions respectively, and model 4 assuming both previous pitch and category information influences current decision-making. The mixed-effect models account for group-level fixed effects and random effects due to variation across participants. The categorical response was modeled using a Bernoulli distribution, *Y*_*ij*_ ∼ *Bernoulli*(*p*_*ij*_), and the full model (model 4) is given by:

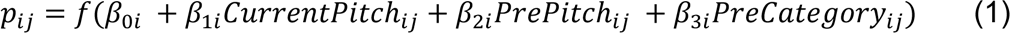

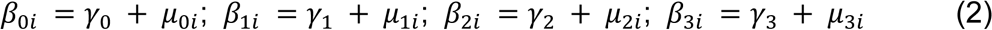

Where *i* indicates participant index and *j* represents the *jth* trial, and 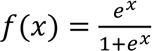 indicates the logistic link function. *Y*_*ij*_ is the categorical report (1 for “high”, 0 for “low”) in the *jth* trial for participant *i*. Parameters *γ*_0_, *γ*_1_, *γ*_2_, *γ*_3_ denote the group-level regression coefficients (fixed effect) for intercept, predictor current pitch, predictor previous pitch, and predictor previous category respectively. The vector μ = (*μ*_0*i*_, *μ*_0*i*_, *μ*_0*i*_, *μ*_0*i*_)^T^ denotes the corresponding individual-level random effect with μ ∼ 𝒩(0, Σ) where Σ is the covariance matrix.

The generalized mixed-effects models were performed using *fitglme* function (Distribution: Binomial, Link: logit) in the Matlab Statistics and Machine Learning Toolbox. Model performance was evaluated using the Akaike information criterion (AIC) with a smaller value indicating better performance. Specifically, we calculated the Δ*AIC* of models 2-4 from the AIC of the model without history influence (model 1).

### EEG data analysis

#### EEG acquisition and pre-processing

The EEG signals were acquired using a 64-electrode actiCAP system and two BrainAmp amplifiers through BrainVision Recorder software (Brain Products). One electrode placed below the right eye was used to record the vertical electrooculography. The impedance of all electrodes was kept below 10 kΩ. During data acquisition, the signals were referenced to electrode FCz, and sampled at 500 Hz. We analyzed the data using Matlab custom codes and the FieldTrip toolbox. Data were epoched 0.5 s before noise onset and 2.25 s after noise onset in Experiment 1, and 0.5 s before response cue and 2 s after noise onset in Experiment 2. The epoched EEG signals were referenced to the mean signal across all electrodes and baseline correction was performed using signals from -0.5 s to -0.3 s relative to noise onset for Experiment 1 and no baseline correction was performed in Experiment 2. Then signals were bandpass filtered between 1 and 30 Hz and down-sampled to 100 Hz. Ocular and other artifacts were removed from the data using independent component analysis (ICA). Guided by visual inspection, epochs with excessive variance were manually excluded from the following analysis. Epochs with reaction times exceeding four standard deviations were also excluded. On average, 40 ± 10 trials in Experiment 1 and 40 + 12 trials in Experiment 2 were excluded.

#### Multivariate pattern analysis

We employed a time-resolved multivariate decoding method ^13, 25^, that is, the Representational Similarity Analysis (RSA)^26^ with Mahalanobis distance^27^ to examine the neural representation of different features (pitch, category, and motor response) across time. At each time point, the neural information linked to each feature was decoded by taking advantage of spatial-temporal voltage dynamics in a predefined time window (20 ms) to increase decoding accuracy. Specifically, at a given time point t, the voltage fluctuations over space (i.e., 64 electrodes) and time (t and t-1) were pooled together as a multivariate pattern (128 values). Using eightfold cross-validation, the Mahalanobis distance between each of the left-out trials and the mean of condition-specific train trials was then calculated with the covariance matrix estimated from the train trials using a shrinkage estimator^53^. Notably, for the seven train-folds, the number of trials for each condition of a certain feature (e.g., pitch) was equalized by randomly subsampling the minimum number of condition-specific trials among different conditions so that the training set would not be biased.

After obtaining the neural representation similarity (Mahalanobis distance) between paired conditions of a certain feature for each trial, we further performed linear regression to quantify the relationship between neural representation similarity and feature similarity. Here, the predictor, feature similarity was dummy coded (0 for pair of the same condition, 1 for pair of different conditions). The averaged regression coefficient of all the trials was used to represent the decoding performance. To obtain reliable estimates of decoding performance, we repeated this procedure 50 times in each of which we randomly divided the dataset into eight folds.

With the same EEG signals, we also computed the decoding performance for each of the current-trial and previous-trial features (pitch, category, and motor responses) independently. Importantly, to exclude potential interference between current and previous features during decoding, we decoded one by fixing the other. For example, when we decoded the pitch information on the current trial, we first divided the trials into five subsets based on the pitches in the previous trials. Then we computed the decoding performance in each subset and then averaged them. Similarly, when decoding the previous-trial pitch, we performed the analysis on trials containing the same current-trial pitch, respectively, and then averaged the results.

#### Statistical analysis

We used a nonparametric sign-permutation test to examine whether the decoding performance was significantly larger than 0 at the group level^54^. For each permutation, the sign of each participant’s decoding performance was randomly flipped with a probability of 50% and the group mean of decoding performance was computed. We conducted permutation 100,000 times and created the null distribution. The group means of original decoding performance were compared against the null distribution and the p-value was calculated as the percentage of values in the null distribution that larger than the real mean. This permutation test was performed for each time point of the whole time course. Then a cluster-based permutation (N = 100,000) was used for multiple comparisons across time in which cluster was defined based on the threshold of p = 0.05. The statistical analysis was performed on the raw data of decoding performance and we smoothed the group means of decoding time course using a Gaussian-weighted moving average filter with a 150 ms time window to better visualize the result when plotting.

#### Cross-correlation analysis

To examine the relative temporal lags of neural representations of different features, we performed cross-correlation analysis for different pairs of decoding performance time series. First, we tested whether the time course differs for different features in the current trial (pitch vs. category, pitch vs. motor response). Next, we investigated whether there was a time lag between the representation of the same feature in previous and current trials.

Specifically, for each paired time series (e..g, current pitch vs. current category), the cross-correlation (xcorr function in Matlab) between their group mean time courses were computed as a function of time lags between the time series. Then we used a nonparametric permutation test^55^ to test its significance. For each permutation, each of the two group mean time courses was shuffled across time following which the cross-correlation was calculated, and then the maximum of the correlation coefficients was selected (multiple comparison corrected). This procedure was conducted 100,000 times to create the null distribution. The permutation test was run to assess whether the original cross-correlation coefficient was higher than the 95% of the values from the null distribution.

#### Neural representation shift analysis

To directly examine the neural representation of serial bias for each feature, we developed a new test to access whether and how the previous-trial feature affects the neural representation of the current-trial feature and in which direction, i.e., attractive or repulsive.

Generally speaking, for each feature (pitch, category, motor response), we built neural templates by averaging the condition-specific trials and then calculated the Mahalanobis distance between each trial and the templates. Specifically, we employed the same 8-fold cross-validation as in the multivariate pattern analysis, that is, first building neural templates by averaging the seven train-folds trials, followed by calculating the Mahalanobis distance between each of the left-out trials and the templates, for each 8-fold division. After repeating this procedure 50 times, we obtained the reliable estimates of neural distance to the templates for each trial. Moreover, considering the strong motor signals in Experiment 1 (Fig. 2A), the motor response was regressed out when performing pitch decoding analysis, to increase the decoding sensitivity. In addition, given the temporal binding between motor response and category in Experiment 2 design (Fig. 5AB), the response cue was controlled when performing category decoding analysis.

We next analyzed whether and how the previous-trial feature influences the neural representation of feature in the current trial.

For pitch neural shift analysis, we took an example of quantifying the influence of past-trial f2 on current-trial f1 for illustration of the general idea (see Figure 3A), as illustrations. First, the neural templates for f1 (dark blue) and f2 (light blue) were selected. Next, we chose trials (current f1, previous f2) and computed the neural distance between each trial in the subsample and the f1 and f2 templates, separately, yielding D1 and D2, from which M_diff (D2-D1) was obtained. For comparison, we chose trials (current f1, previous f1) as baselines and did the same distance computation, yielding the M_Diff (D2-D1). Finally, the difference between M_diff and M_Diff was calculated (Shift_dist) to quantify the influence of the previous f2 on the neural representation of current f1. If previous f2 attracts f1 neural representation, we would expect smaller M_diff than M_Diff, yielding positive Shift_dist values, and vice versa. There are three possibilities – attractive, no effect, and repulsive (Fig. 3A, right panel), corresponding to positive, around zero, and negative Shift_dist values, respectively. The neural shift analysis was performed at each time point, for each subject.

For category neural shift analysis (see supplementary Fig. 2), we first built the neural templates for the “High-pitch” (dark green circle) and “Low-pitch” (light green circle) categories, based on all trials (approximated by eight-fold cross validation as the above Pitch shift analysis). Next, we computed the neural distance between each trial and the “High” and “Low” templates, yielding D1 and D2, respectively. Positive serial bias would predict neural attraction to the past-trial category, that is, positive Diff_High (D1-D2) or positive Diff_Low (D2-D1) values when preceded by the “High” and “Low” category, correspondingly. We then averaged the Diff-High and Diff_low values as Shift_dist to characterize the neural shift of category information.

There are three possibilities – attractive, no effect, and repulsive, corresponding to positive, around zero, and negative Shift_dist values, respectively. The neural shift analysis was performed at each time point, and in each subject.

A similar idea as category analysis has been applied to motor response. Specifically, we first built the neural templates for the “Left” and “Right” motor responses, based on all trials. Next, we computed the neural distance between each trial and the “Left” and “Right” templates, yielding D1 and D2, respectively. Positive serial bias would predict neural attraction to the past-trial category, that is, positive Diff_High (D1-D2) or positive Diff_Low (D2-D1) values when preceded by the “Left” and “Right” categories, correspondingly.

We then averaged the Diff-High and Diff_low values as Shift_dist to characterize the neural shift of category information. There are three possibilities – attractive, no effect, and repulsive, corresponding to positive, around zero, and negative Shift_dist values, respectively. The neural shift analysis was performed at each time point and in each subject.

A similar nonparametric sign-permutation test and cluster-based multiple comparison correction were performed on the neural shift analysis results.

Finally, to examine the behavioral relevance of the neural shifts, for each feature, we divided participants into two groups (high-bias vs. Low-bias) based on behavioral serial bias (the regression coefficient extracted from the GLM model 4) and then performed the same neural shift analysis for the two groups, respectively.

## Acknowledgments

This work was supported by the National Science and Technology Innovation 2030 Major Program 2021ZD0204103 to H.L., National Natural Science Foundation of China (31930052) to H.L, and China Postdoctoral Science Foundation (2020M680166) to H.Z. We thank Dr. Nai Ding, Dr. Jian Li for their helpful comments.

## Notes

### Competing Interest Statement

The authors have declared no competing interest.

