## Supplementary materials for "Co-occurrence of past and present shifts current neural representations and mediates serial biases"

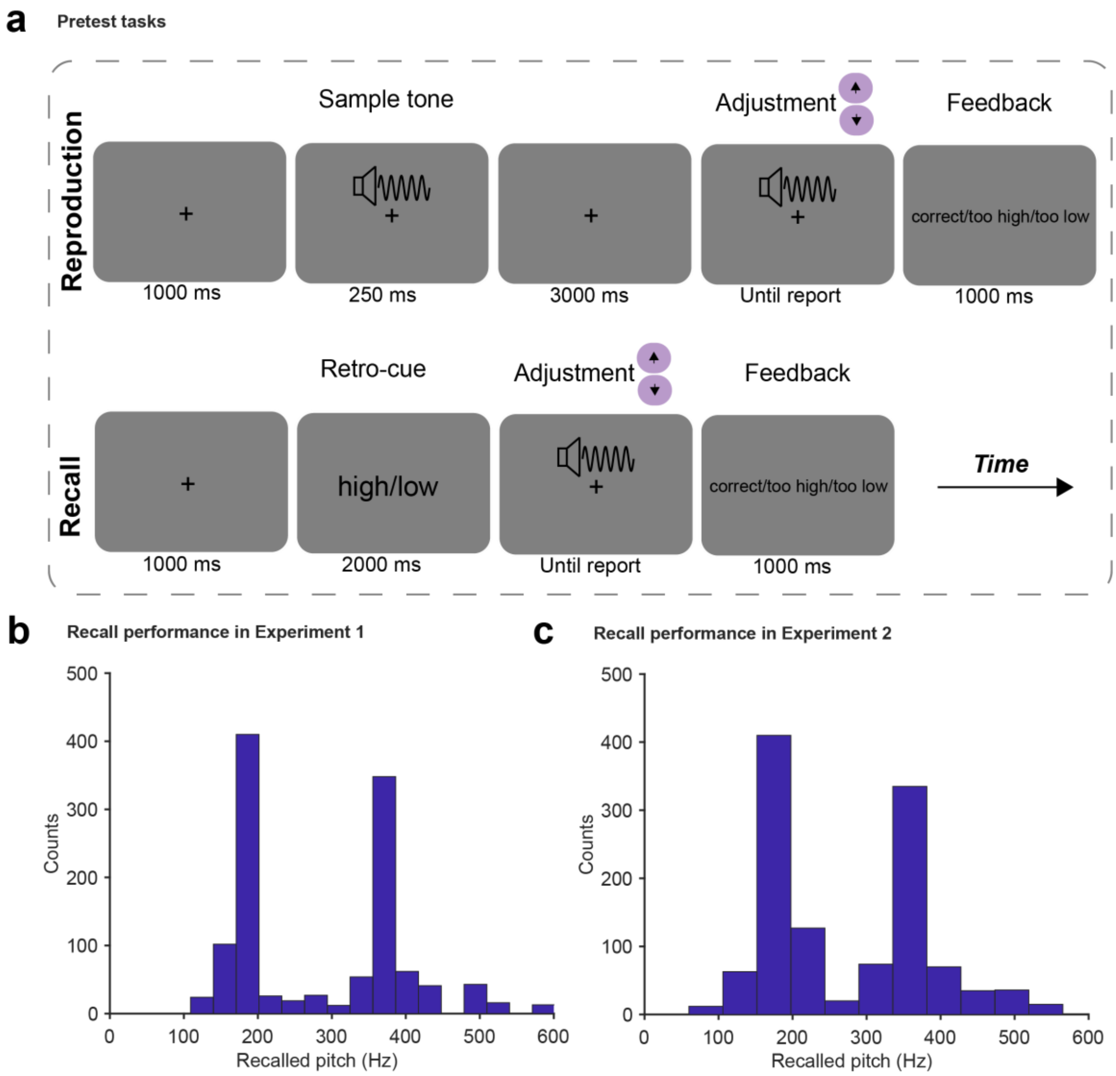


**Supplementary Figure 1. Pretest task paradigm and behavioral performance. A**. Tone memory task paradigm. Upper: A reproduction task where subjects were required to reproduce the given tone (180 or 360 Hz) by using the up and down arrow keys, after which feedbacks were provided. Lower: A recall task where subjects had to recall the respective tone according to the retro-cue “high” (360 Hz) or “low” (180 Hz) and reproduce it. The pretest contains a block-design session (two reproduction blocks and two recall blocks each of which testing only one tone for 10 trials) and a randomization design session (one reproduction block and one recall block each of which testing two tones in a trial-by-trial random manner, 20 trials for each tone). **BC**. Recall performance in the randomization design session of Experiments 1(B) and 2 (C). Both plots show two clear peaks around 180 Hz and 360 Hz (aggregate results across subjects).

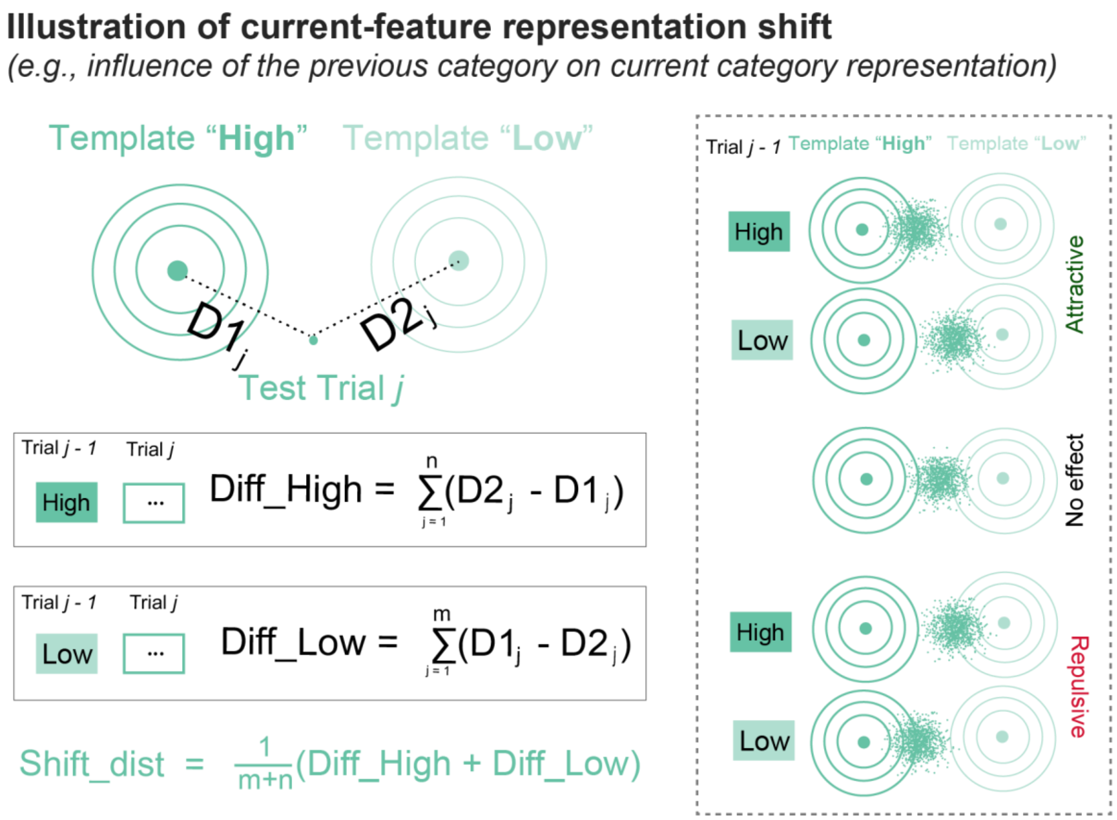


**Supplementary Figure 2. Illustration of neural representational shift analysis (category as an example). Left**: two neural templates were built for "High" (dark green circle) and "Low" (light green circle) category base on all trials. For each trial, its neural distance to the two templates were computed, resulting in D1 and D2, respectively. Positive serial bias would predict neural attraction to previous category, i.e., positive Diff_High (D2-D1) and positive Diff_Low (D1-D2) values when preceded by "High" and "Low" category, respectively. The two values were averaged as Shift_dist to characterize the neural shift for Category. **Right**: Neural representation of current-trial category is attracted toward (upper, positive Shift_dist values), repulsed from (lower, negative Shift_dist values), or not affected (middle, around zero Shift_dist values) by prior category information.

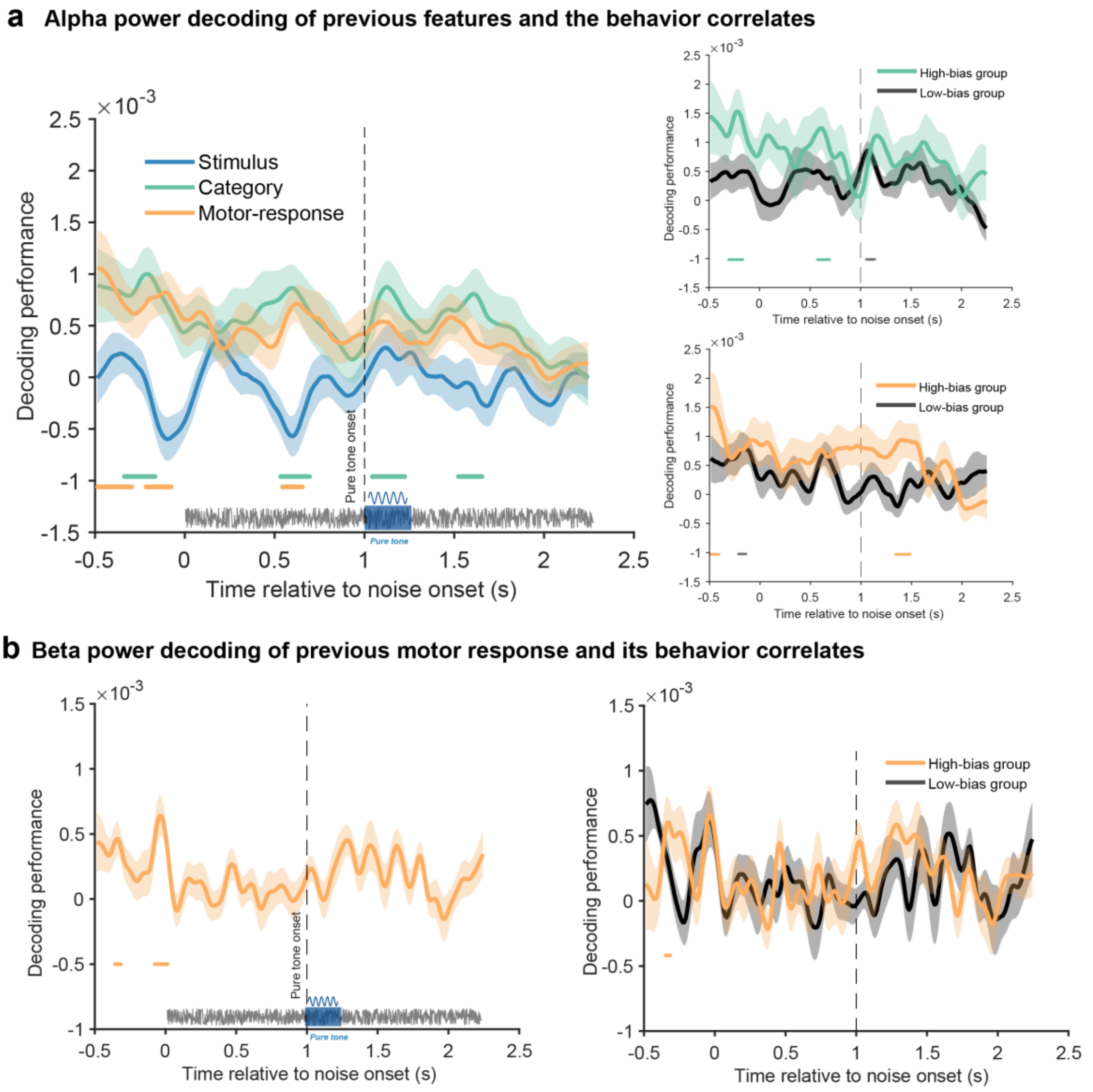


**Supplementary Figure 3. Alpha and beta power decoding performance of previous features and the behavioral relevance. A.** Alpha power decoding performance of previous features and the behavioral relevance. Left: Grand average alpha power decoding performance for previous features as a function of time following white noise onset (vertical dotted line), for Pitch (blue), Category report (green), and Motor response (orange). The vertical dashed line denotes the tone onset. Horizontal colored lines denote significant temporal clusters (cluster-based permutation test, one-sided, corrected, $p < 0.05$) for each feature. Shadows represent SEM. Right: Subjects were divided into two groups based on the serial bias behavior in Category, or Motor response, respectively. Grand average alpha power decoding performance of High-bias (colored lines) and Low-bias (black line) groups for Category (Upper), and Motor response (Lower). Horizontal lines denote significant temporal clusters (cluster-based permutation test, one-sided, corrected, $p < 0.05$). **B.** Beta power decoding performance of previous motor response and the behavioral relevance. Left: Grand average beta power decoding performance for previous motor response as a function of time following white noise onset. Right: Grand average beta power decoding performance of High-bias (orange line)

and Low-bias (black line) groups. Horizontal lines denote significant temporal clusters (cluster-based permutation test, one-sided, corrected, $p < 0.05$).
